# Generating Functional Cells Through Enhanced Interspecies Chimerism with Human Pluripotent Stem Cells

**DOI:** 10.1101/2022.02.10.479858

**Authors:** Yanling Zhu, Zhishuai Zhang, Nana Fan, Ke Huang, Hao Li, Jiaming Gu, Quanjun Zhang, Zhen Ouyang, Tian Zhang, Jun Tang, Yanqi Zhang, Yangyang Suo, Chengdan Lai, Jiaowei Wang, Junwei Wang, Yongli Shan, Mingquan Wang, Qianyu Chen, Tiancheng Zhou, Liangxue Lai, Guangjin Pan

## Abstract

Obtaining functional human cells through interspecies chimerism with human pluripotent stem cells (hPSCs) remains unsuccessful due to its extreamly low efficiency. Here, we show that hPSCs failed to differentiate and contribute teratoma in presence of mouse PSCs (mPSCs). While, MYCN, a pro-growth factor dramatically promotes hPSC contribution in teratoma co-formation by hPSCs/mPSCs. MYCN combined with BCL2 (M/B) greatly enhanced conventional hPSCs to integrate into preimplantation embryos of different species, such as mice, rabbits and pigs and substrantially contributed to mouse post-implantation chimera in embryonic and extra-embyonic tissues. Strikingly, M/B-hPSCs injected into preimplantation *Flk-1*^+/-^ mouse embryos show further enhanced chimerism that allows obtaining live human CD34^+^ blood progenitor cells from chimeras through cell sorting. The chimera derived human CD34^+^ cells further gave rise to various subtype blood cells in typical colony forming unit(CFU) assay. Thus, we proof the concept to obtain functional human cells through enchanced interspecies chimerism with hPSCs.

## Introduction

Interspecies chimerism using human pluripotent stem cells (hPSCs) is a promising approach for the generation of xenogenic organs through animal blastocyst complementation^1-5^. The stragegy that *Pdx1*-deficient murine blastocysts injected with wild-type rat PSCs has been shown to generate fully functional pancreas tissue of rat origin^1^. However, hPSCs exhibit very limited efficiency in interspecies chimerism even though they can self-renew and are pluripotent, generating a large variety of cell types upon differentiation *in vitro* and teratoma formation *in vivo*^4,6-8^. One explanation for this limited efficiency is that conventional hPSCs are in a primed state and thus do not match the developmental stage of the preimplantation blastocyst^3,9-12^. To overcome this barrier, intensive efforts have been made to generate naïve hPSCs with cellular and transcriptional properties similar to those of naïve pluripotent cells^13-20^. However, these naïve hPSCs generated by different protocols are highly variable in terms of the effciency in interspecies chimerism^7,13,21,22^, indicating that additional barriers remain to be illuminated. Indeed, we have shown that the conventional hPSCs undergo rapid apoptosis when injected into preimplantation blastocysts in part due to activation of the *Ink4a* pathway^23,24^. Consistently, overexpression of BMI1, which suppresses *Ink4a* or BCL2 to directly inhibit apoptosis, enables primed hPSCs to integrate into the early embryos of mice, rabbits and pigs and to differentiate into both embryonic and extraembryonic cell types in mouse chimeras^24,25^. Recently, Wu et al. reported a cell competition between hPSCs and host cells in interspecies chimerism, which leads to hPSC apopotosis during chimera^26^. However, eventhough the apoptosis is blocked in hPSCs by anti-apoptosic factor such as BCL2 or BMI1, the chimera efficiency is still extreamly low and obtaining live and functional hPSCs derived cells in chimera remains unsuccessful.

To investigate new strategies that enhance hPSC interspecies chimeasim, we firstly set up an teratoma co-formation assay by mixed hPSCs and mouse ESCs (mPSCs), which to some extent mimicks chimara development *in vivo*. We showed that hPSCs failed to co-differentiate with mESCs and contribute teratoma in the presence of mESCs. Overcoming apoptosis by BCL2 in hPSCs promoted their contribution rate from 0.5% to 20% in teratomas formed by a mixture with mESCs at the original 1:1 ratio. Combined with MYCN, a pro-growth factor (M/B) promoted hPSC contribution rate to 75% in teratoma co-formed by hPSCs/mESCs. M/B-hPSCs show robust integration into preimplantation embryos of different species, such as mice, rabbits and pigs. Strikingly, M/B-hPSCs injected into preimplantation *Flk-1*^+/-^ mouse embryos show further enhanced chimera efficiency that allows obtaining live hPSCs derived CD34^+^ blood progenitor cells from chimeras through cell sorting. Thus, we proof the concept to obtain live and functional hPSC derived cells through interspecies chimerism.

## Results

### Promoting contribution of hPSCs in teratomas co-formed by hPSCs/mESCs

The embryonic development timing is distinct between human and other specises, such as mouse. To exmine whether hPSCs and mESCs could differentiate normally in the presence of each other, we mixed differentially labeled hPSCs and mESCs in a 1:1 ratio for teratoma co-formation (**Fig. 1A-C**). The hPSC/mESC mixture formed typical teratomas after injection into immunodeficient mice (**Fig.1B**). However, the numbers of DsRed-labeled cells derived from hPSCs were very low in co-formed teratomas, accounting for fewer than 1% of all cells. While, the majority of cells were GFP-labeled mESCs derived cells (**Fig. 1B-C**), indicating hPSCs failed to differentiate normally in the presence of mESCs. Expressing BCL2 in hPSCs (**Fig.1E and Fig.S1**) to overcome potential apopotosis during teratoma formation only promote their contribution rate up to around 20% in co-formed teratoma (**Fig. 1F-H**). To search additional factors to promote hPSC co-differentiation with mPSCs, we focus on MYC family genes, the well-known factors to promote cell growth. *MYCN*, while not *C-MYC* showed relatively higher expressions in mESCs compared with hPSCs (**Fig. 1D**). Strikingly, *BCL2* combined with *MYCN* expression (M/B) promoted hPSC contribution rate up to 70% in teratoma formed by hPSCs/mESCs (**Fig. 1E-H and Fig. S1**). Notably, immunohistochemical analysis and hematoxylin and eosin (HE) staining showed three typical germ layers in teratomas formed by the mESC/B- or M/B-hPSC mixture (**Fig. 1H-I**). Together, these data demonstrate that BCL2 combined with MYCN enables enhanced differentaiton of hPSCs in the presence of mESCs in teratoma co-formation assay that might to some extent mimick chimera development *in vivo*.

**Figure 1.**
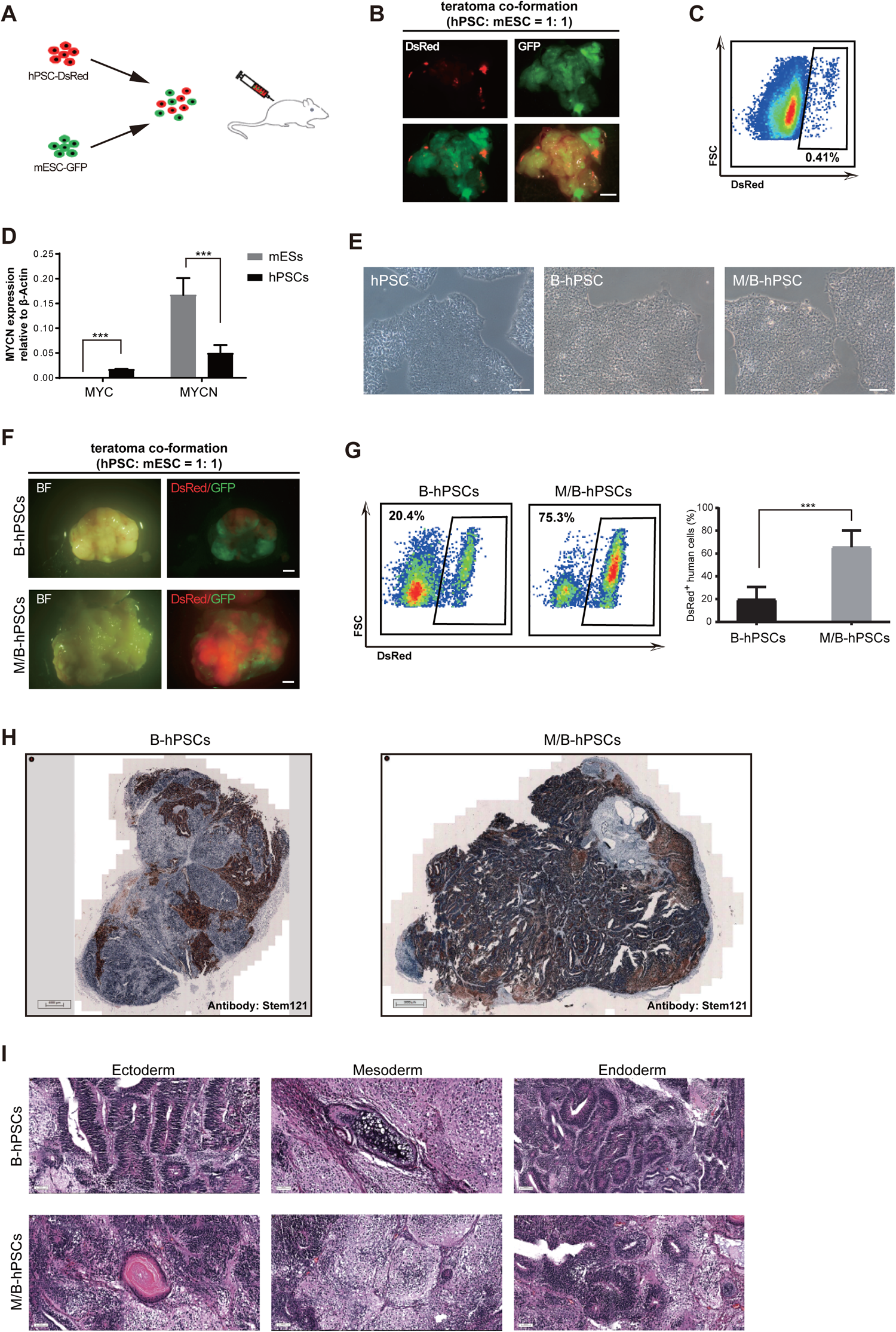
MYCN/BCL2 promotes the contribution of hPSCs in teratomas co-formation mixed with mESCs. **(A)** Schematic overview of co-differentiation of hPSCs-DsRed and mESCs-GFP in vivo. hPSCs-DsRed and mESCs-GFP are initially mixed in 1:1 ratio at single cell for further differentiation. **(B-C)** The teratoma formed by the mixture of hPSCs-DsRed and mESCs-GFP at four weeks. Dsred labeled human cells in teratoma were analyzed by flow cytometry. Scale bars, 2 mm. **(D)** Analysis of the MYC and MYCN expression in the indicated cells by RT-qPCR. Error bars represent mean+SEM of three parallel experiments. ***p < 0.001. **(E)** The morphology of BCL2 or MYCN/BCL2 expressed hPSCs. Scale bars, 100 µm. **(F)** Teratomas formed by B-hPSCs or M/B-hPSCs mixed with mESCs in 1:1 ratio. Scale bars, 2 mm. **(G)** Flow cytometry analysis of Dsred labeled human cells in indicated teratomas. Error bars represent mean+SEM of three independent replicates. ***p < 0.001. **(H)** Immunohistochemical analysis of human cells in indicated teratomas with human specific anti-Stem121 antibody. Scale bars, 1 mm. **(I)** H&E staining of teratomas formed by B-hPSCs or M/B-hPSCs mixed with mESCs in 1:1 ratio. Three typical germ layers are shown. Scale bars, 50 µm.

### M/B-hPSCs show enhanced integration and contribution in interspecies pre-implantation blastocysts

We and others have shown that primed hPSCs expressing antiapoptotic factors such as BMI1 and BCL2 contribute to both extraembryonic and embryonic lineages when injected into interspecies pre-implantation blastocysts^24,25^, but the overall efficiency is quite limited. To examine whether combination of MYCN/BCL2 can further promote interspecies chimerism, we injected hPSCs expressing BCL2 (B-hPSCs) or hPSCs expressing both BCL2 and MYCN (M/B-hPSCs) into 8-cell (8C)-stage mouse, pig or rabbit embryos (**Fig. 2A**). The integration of DsRed-labeled human cells was examined after 48-60 hours of culture *in vitro*. M/B-hPSCs showed significantly higher cell numbers in mouse, pig and rabbit embryos than B-hPSCs (**Fig. 2A-B**).

**Figure 2.**
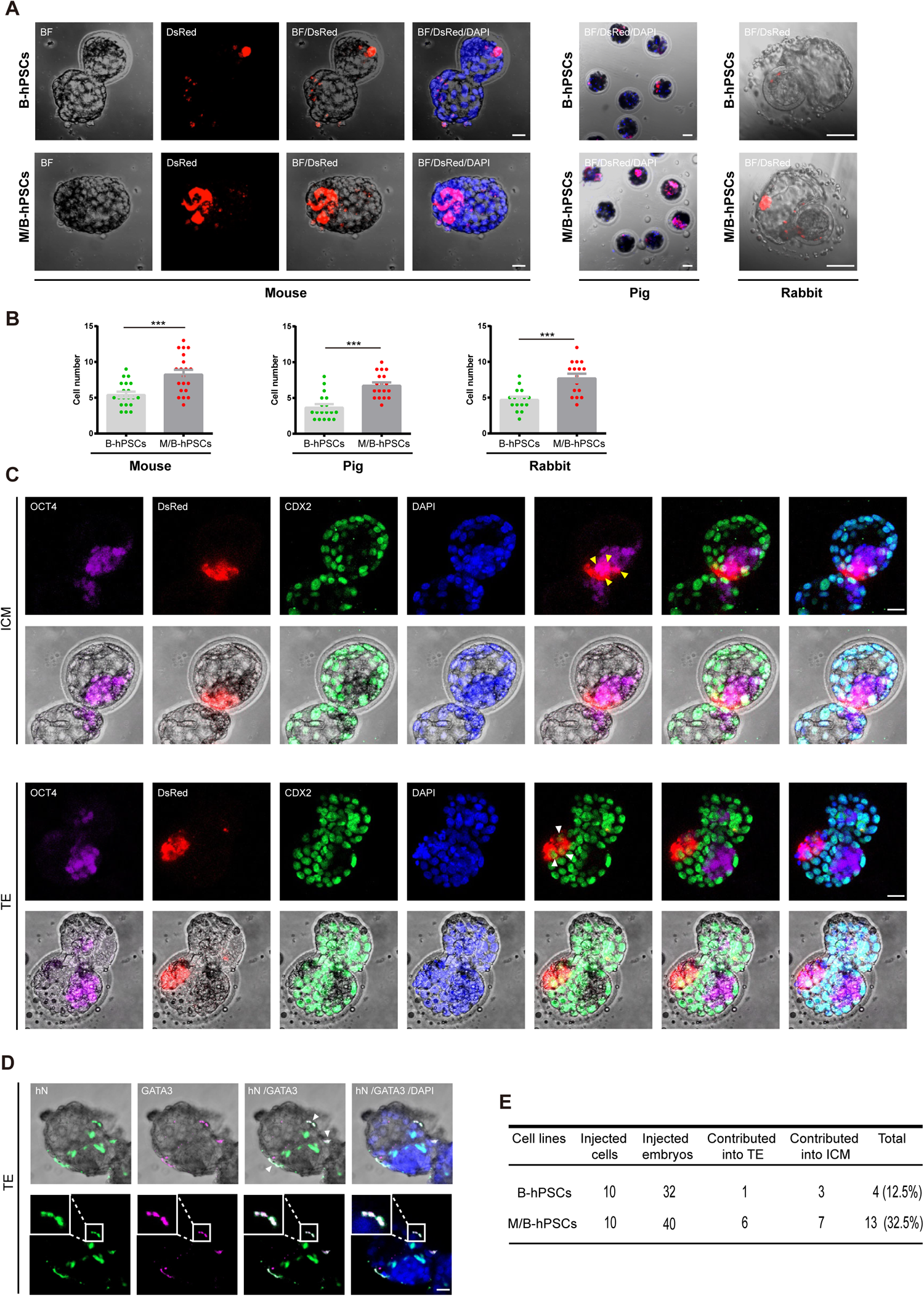
MYCN/BCL2 promotes the integration and proliferation of hPSCs in inter-species pre-implantation embryos. **(A)-(B)** Survival and proliferation of Dsred labeled hPSCs in pre-implantation embryos of different species. DsRed+ hPSCs (10 cells for mouse, 3-5 cells for pig and 7 cells for rabbit) were injected into 8-cell (8C) stage embryos of indicated species and cultured for 3 days in vitro and the cell number of each indicated cell lines was calculated. Scale bars, 20 µm. Mouse: mean + SEM of 20 (+B) or 20 (+N+B) samples; Pig: mean + SEM of 18 (+B) or 18 (+N+B) samples; Rabbit: mean + SEM of 16 (+B) or 16 (+N+B) samples. ***p < 0.001. **(C)** Contribution of hPSCs to embryonic and extra-embryonic embryonic lineages in mouse chimeras. Representative images showing the integrated of M/B-hPSCs co-express OCT4, ICM marker or CDX2, early trophoblast marker in cultured chimeric embryos. Yellow arrow, DsRed^+^/OCT4^+^ cells; white arrow, DsRed^+^/CDX2^+^ cells; scale bars, 20µm. **(D)** Representative images showing the integrated of M/B-hPSCs co-staining with Hn (human cell nucleus specific antibody) and GATA3 (early extra-embryonic lineages marker) in cultured chimeric embryos. white arrow, hN^+^/GATA3^+^ cells; scale bars, 20µm. **(E)** Summary of chimera assays with injection of ten indicated DsRed+ cells at the 8C stage embryo, and followed 48–60h in vitro development into blastocyst stage.

We then examined the cell fate of these integrated hPSCs injected in mouse pre-implantation embryos. The extraembryonic tissues (ExEms) as well as the inner cell mass (ICM) were then examined by coimmunostaining for the trophoblast marker CDX2 or GATA3 and the ICM marker OCT4 (**Fig. 2C**). A considerable number of M/B-hPSCs were detected as CDX2/GATA3- or OCT4-positive injected in mouse embryos, indicating that they contributed to both extraembryonic and embryonic lineages (**Fig. 2C-D**). Moreover, M/B-hPSCs showed significantly higher efficiencies than B-hPSCs in terms of their lineage contributions in pre-implantation mouse blastocysts (**Fig. 2E**). Overall, these data demonstrate that the combination of MYCN/BCL2 promotes interspecies chimerism of hPSCs in pre-implantation blastocysts of different species.

### M/B-hPSCs show increased integration in mouse post-implantation chimera

We then examined the long-term chimeric contributions of M/B-hPSCs with in mouse post-implantation E10.5 conceptuses. First, by using a widely used human mitochondrial DNA (hmtDNA) qPCR assay with 1/10^4^ as a threshold, 70.8% of mouse embryos injected with M/B-hPSCs contained human cells, while only 21.1% of mouse embryos injected with B-hPSCs contained human cells (**Fig. 3A-B**). Moreover, M/B-hPSCs contribution were detected in both extra- and embryonic lineages in recovered embryos (**Fig. 3A**). In contrast, it’s hard to detect B-hPSCs in contributing to both extra- and embryonic lineages in E10.5 chimeras (**Fig. 3B**). To confirm the precence of human cells in chimera, we then examined human cells through immunostaining. We detected a considerable number of human cells at various embryonic regions in E10.5 mouse chimeras injected with DsRed-labeled M/B-hPSCs by immunostaining with the human-specific antibody Stem121 (**Fig. 3 C and Fig. S4D**). These cells differentiated into various morphologically distinct lineages at different embryonic regions in the mouse chimeras (**Fig. 3C and Fig. S4D**). Furthermore, the DsRed+ cells in placenta tissue could be co-stained with antibody against CK7, a trophoblast markers. (**Fig. S4E**). In all, these data demonstrate that the combination of MYCN/BCL2 largely promotes the long-term integration of hPSCs in mouse post-implantation embryos.

**Figure 3.**
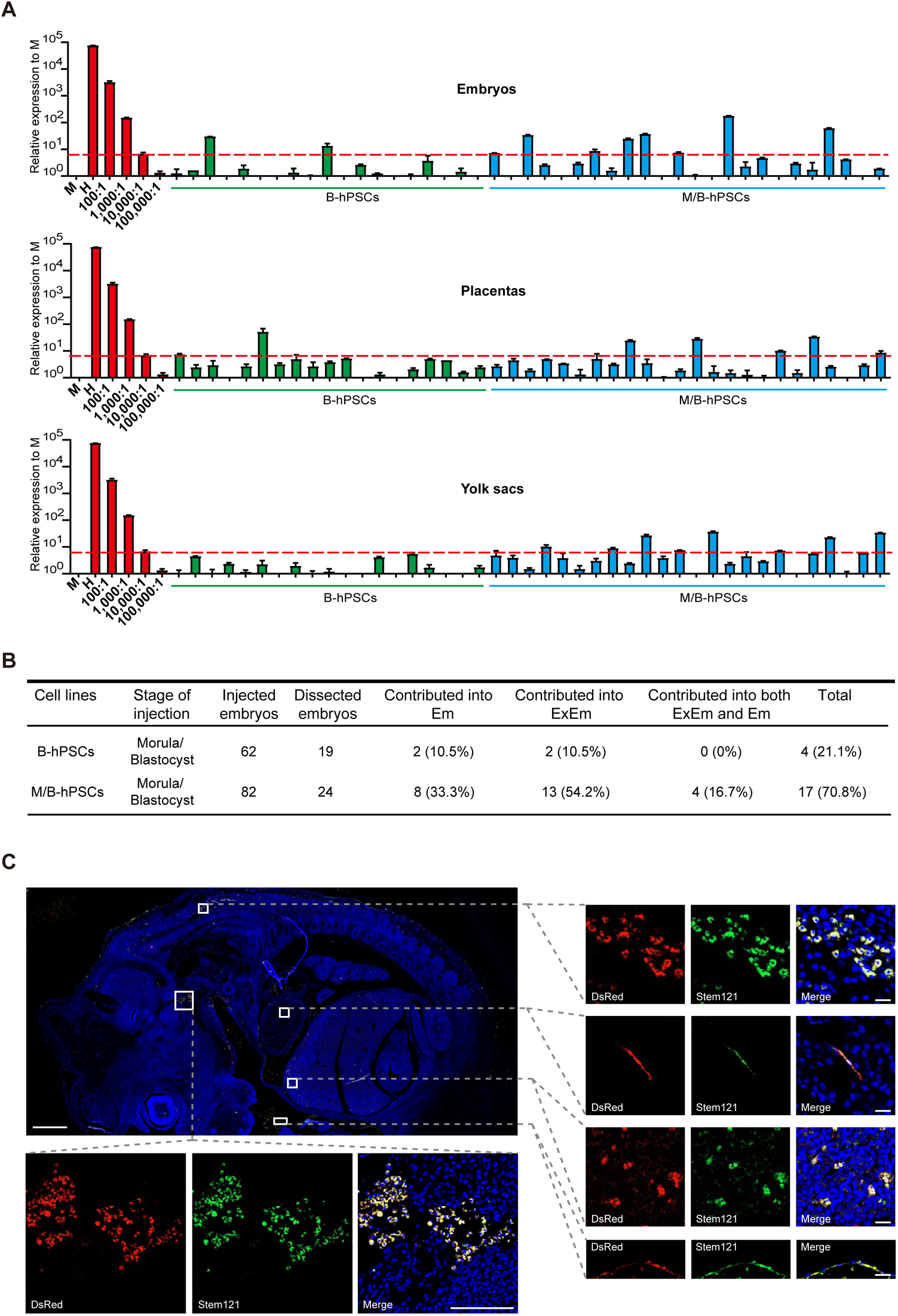
MYCN/BCL2 promotes long-term chimera of hPSCs in post-implantation mouse embryos. **(A)** Quantitative genomic PCR analysis of the human mitochondria DNA in E10.5 chimeras of mouse embryos, placentas and yolk sacs injected with the indicated cells. A human DNA control (H), a mouse DNA control (M) and a series of human-mouse cell dilutions (1/100 to 1/100,000) were run in parallel to estimate the degree of human cell integration. The dashed line indicates the detection level of one human cell in 10,000 mouse cells. Error bars represent mean+SEM of three parallel experiments. **(B)** Summary of mouse chimeras containing human cells. Embryos were recovered at E10.5 stage. ExEm: extra-embryonic tissues including both placentas and yolk sacs, Em: embryonic lineages. **(C)** Analyses of human cells at different embryonic regions in E10.5 mouse chimera by immunofluorescence using human specific marker Stem121. Scale bars, 500 µm (D-left), 20 µm (D-middle), 5 µm (D-right), 500 µm (E-left), 100 µm (E-right).

### M/B-hPSCs generate blood progenitor cells in *Flk-1* haplodeficient mouse embryos

Interspecies complementation using hPSCs is considered a promising approach to generate humanized functional cells or organs. During interspecies complementation, the injected PSCs are required to complement the deficient lineages in host embryos, which might in turn enhance their overall contribution rate in chimera. We then examined whether M/B-hPSCs could generate targeted cells **i**n *Flk-1* impared mouse embryos, which have severe defects in blood/endothelia lineages^27,28^. In this model, the mouse *Flk-1* gene locus was replaced with EGFP (*Flk-1*^+/EGFP^)^28^. DsRed-labeled M/B-hPSCs were injected into pre-implantation blastocysts that were generated by intercrossing *Flk-1*^+/EGFP^ mice and analyzed at E10.5. Since the homozygous *Flk-1*^EGFP/EGFP^ embryos failed to develop to E10.5, we just obtained and analyzed chimeras in *Flk-1*^+/EGFP^ embryos in which the mouse endothelial lineage was labeled by EGFP (**Fig. 4A**)^28^. M/B-hPSC-derived cells were detectable in nearly 90% of *Flk-1*^+/EGFP^ mouse chimeras (**Fig. 4B-C**). Moreover, the overall contribution of human cells was siginificantly higher in *Flk-1*^+/EGFP^ mouse chimeras compared with WT chimeras **(Fig. 3 and Fig. 4B**). In some chimeras, the human cell ratios were above 1:1000 or even 1:100 based on an hmtDNA qPCR assay (**Fig. 4B**). Importantly, a number of live human CD34^+^ hematopoietic/endothelial progenitor cells were detected in *Flk-1*^+/EGFP^ mouse chimeras based on FACS analysis (**Fig. 4D**). These cells were successfully sorted and cultured *in vitro* (**Fig. 4E**). To examine the function of these chimera derived human CD34^+^ cells, we performed a colony-forming unit (CFU) assay (**Fig. 4F**). Strikingly, CD34^+^ human cells sorted from the chimeras successfully generated typical CFUs *in vitro* (**Fig. 4F**). Further Giemsa staining of cells from these colonies showed morphologies of various blood lineages, such as macrophage, erythroid, monocyte, granulocyte, and other lineages (**Fig. 4G**). Together, these data demonstrate that the chimersim enhanced hPSCs generate functional blood progenitor cells in *Flk-1* haplodeficient mouse embryos through complementation.

**Figure 4.**
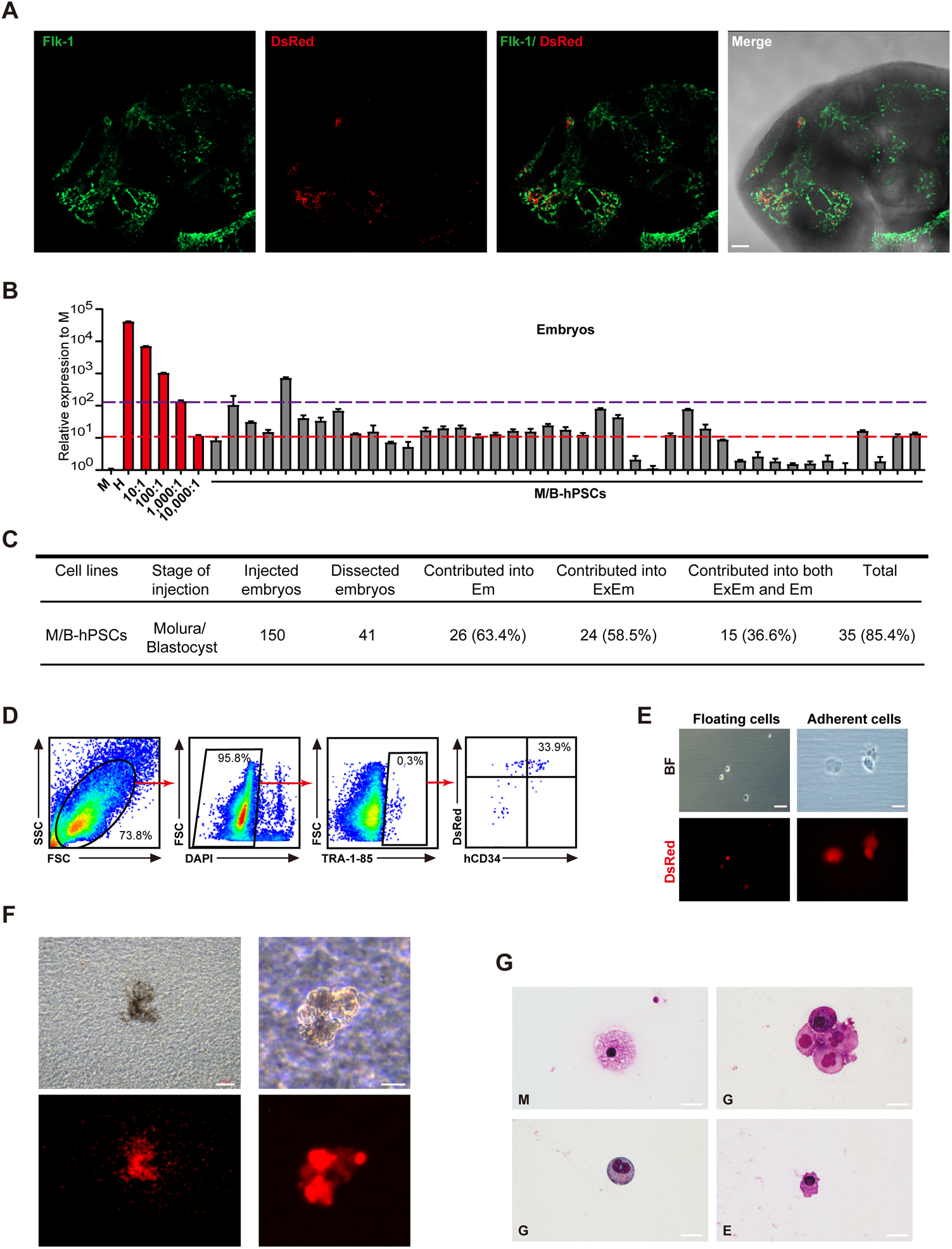
Generation of human CD34^+^ endothelial/blood progenitor cells through inter-species chimera. **(A)** Representative fluorescence images showing the integration of human cells in the vascular regions of E10.5 *Flk-1*^*+/EGFP*^ mouse chimeric embryos. Scale bars, 100 µm. **(B)** quantitative genomic PCR analysis of the human mitochondria DNA in E10.5 *Flk-1*^*+/EGFP*^ mouse chimeras injected with M/B-hPSCs. A human DNA control (H), a mouse DNA control (M) and a series of human-mouse cell dilutions (1/100 to 1/100,000) were run in parallel to estimate the degree of human cell integration. The red and purple dashed lines indicate the detection level of one human cell in 10,000 and 1,000 mouse cells, respectively. Error bars represent mean+SEM of three parallel experiments. **(C)** Summary of mouse chimeras containing human cells. Embryos were recovered and analyzed at E10.5 stage. ExEm: extra-embryonic tissues, including both placentas and yolk sacs, Em: embryonic lineages. **(D)** Representative flow cytometry analysis of live human cells in E10.5 *Flk-1*^*+/EGFP*^ mouse chimeric embryos. The antibody TRA-1-85 was used to identify human cells, hCD34 was used to identify human endothelial and blood cells. **(E)** The morphology of sorted live human cells (TRA-1-85^+^). Scale bars, 100 µm. **(F)** Representative pictures of CFUs formed by the sorted live human cells (TRA-1-85^+^). Scale bars, 200 µm (left), 100 µm (right). **(G)** May–Grunwald–Giemsa staining of different blood cells from (F). Scale bars, 20 µm. E, erythroid; G, granulocytes; M, macrophages.

## Discussion

Interspecies chimerism using hPSCs is hoped to generate xenogenic organs or functional cells through animal blastocyst complementation. However, the concept has not been proved due to the extremely low efficiency of hPSCs in interspecies chimera. Even the “stage-matched” naïve hPSCs still show very low and variable interspecies chimera competence^7,13,21,22^. In this study, we generated a chimerism enchanced hPSCs by combinating anti-apoptotic factor, BCL2 and pro-growth factor MYCN (M/B). M/B expression greatly promoted integration and growth of conventional hPSCs in pre-implantation blastocysts of different species such as mouse, rabbit and pig as well as in post-implantation of mouse embyos. The chimera efficiency was further increased in *Flk-1* haplodeficient mouse embryos, indicating the context of complementation might enchance chimera formation. Importantly, we could sort the live human CD34^+^ blood/endothelial progenitor cells from mouse chimera with M/B-hPSCs. These chimera derived human CD34^+^ progenitors further gave rise to various subtype blood cells in CFU assay. To our knowledge, obtaiing live and functional human cells from inter-species chimera has not been reported. Thus, our study proves the concept to generate functional human cells through enhanced interspecies chimerism with hPSCs.

The mechanism of chimera barrier using hPSCs remains to be fully illustrated. We and others have reported that hPSCs undergo severe apoptosis after injection into animal pre-implantation blastocysts^23-25^. Recently, Wu et al. reported a cell competition mechanism between hPSCs and host cells, leading the death of hPSCs in chimera^26^. Indeed, hPSCs contributed very few cells in teratoma co-formation by the mixture containing mESCs (**Fig. 1**). The failure of hPSCs to differentiate in the presence of mESCs might not due to their “primed state”, since hPSCs also underwent severe apoptosis in the presence of mouse primed epiblast stem cells (mEpiSCs)(Fig. S4A). The anti-apoptotic factor BCL2 did elevate the ratio of hPSCs derived cells in teratoma, but still was much lower compared with mESCs derived cells (**Fig. 1**). Interestingly, combining with antother pro-growth factor, MYCN enables hPSCs dominating the teratoma co-formation with mESCs (**Fig. 1**). Finally, BCL2 and MYCN successfully enhanced interspecies chimera in hPSCs, as observed in co-teratoma formation. Interestingly, MYCN factor was also reported to enhance cell competition in mouse early embryonic development, during which MYCN-high epiblast cells elimated MYCN-low ones^29^. hPSCs indeed show relatively lower MYCN expressions compared with mESCs (**Fig. 1D**), thus might be eliminated during co-teratoma or chimera formation. Indeed, in an simple co-differentaiton assay mixed by hPSCs and mESCs, hPSCs underwent rapid apoptosis and was eliminated (**Fig. S2 and Fig. S4F**). While, MYCN expresson largely prevent cell death and promote the growth of hPSCs during co-differention with mESCs (**Fig. S3A-D**). hPSCs expressing MYCN or both BCL2 and MCYN exibit normal morphology and differentiation potential in forming typical teratoma with normal three germ layers (**Fig. S1**). Notablly, BCL2 and MCYN could enable hPSCs to survival well in co-differentiation or teratoma formation mixed with mESCs in a very low ratio such as 1:30 (Fig. S4B-C). However, just MYCN itself did not overcome apoptosis or enhance cloning efficiency in dissociated hPSCs, as well as failed to promote hPSC intergration in pig pre-implantation embyos (**Fig. S3E-F**). Thus, combining BCL2 and MYCN provides an enhanced approach for interspecies chimersim using hPSCs, which marks a important progress by obtaining live and functional human cells through interspecies chimerism.

## Acknowledgements

We thank the lab members in GIBH for their kindly help. This work was supported by Strategic Priority Research Program of Chinese Academy of Sciences (Grant No. XDA16030504); Guangzhou Science and Technology Program General project (201904020045); the National Natural Science Foundation of China (31801220, 31971374, 32071454); the National Key Research and Development Program of China, Stem Cell and Translational Research (2017YFA0102600); China Postdoctoral Science Foundation (2020M672589, 2020T130134); Science and Technology Planning Project of Guangdong Province, China (2017B030314056); the Frontier and Key Technology Innovation Special Grant from the Department of Science and Technology of Guangdong Province (2016B030230002, 2016B030229008); Key Research & Development Program of Bioland Laboratory (Guangzhou Regenerative Medicine and Health Guangdong Laboratory) (2018GZR110104005); the Guangdong Province Special Program for Outstanding Talents (to G.P., 2019JC05Y463).

## Author contributions

G.P., L.L. and Y.Z. designed the project and analyzed the data. G.P., Y.Z. and Z.Z. wrote the manuscript. Y.Z., Z.Z. and N.F. performed most experiments and analyzed the data. Z.Z., K.H., H.L., J.G., Q.Z., Z.O., Y.S., C.L., and J.W. performed the animal experiments. T.Z., J. T. and J.W. assisted with the FACS. J.T. and Y.Z. performed the qPCR assays. T.Z. performed karyotype analysis. Y.S. and Q.C. gave experiment suggestions or provided experiment materials for this research. All authors read and approved the final manuscript.

## Methods

### Culture and maintenance of hPSCs and mESCs

hPSCs were cultured under 20% O_2_ and 5% CO_2_ at 37 °C condition. The plates should be coated by Matrigel before use. Human PSC lines UH10-DsRed, UH10-DsRed+MYCN, UH10-DsRed+BCl2 and UH10-DsRed+MYCN+BCl2 were maintained on plates with hPSCs medium mTeSR1 (STEM CELL). Every 3 days, the human PSC lines were passaged using 0.5mM ethylenediaminetetraacetic acid disodium salt (EDTA-2Na). All of the cell lines indicated above have been tested to be free of mycoplasma contamination. In particular, the generation of the UH10 hiPSCs was approved by the Institutional Review Board at Guangzhou Institutes of Biomedicine and Health. Also, we have complied with all relevant ethical regulations and obtained consents from human participants.

mESCs were cultured under 20% O_2_ and 5% CO_2_ at 37 °C condition. The plates should be coated by gelatin before use. mESCs were maintained in N2B27 + 2iL medium (50% DMEM/High glucose [Hyclone], 50% Knockout DMEM [Gibco], N2 [Gibco, 200×] + B27 [Gibco, 100×], NEAA [Gibco, 100×], GlutaMAX [Gibco, 100×], Sodium Pyruvate [Gibco, 100×], 1 μM PD0325901 [Selleck], 3 μM CHIR99021 [Selleck], 100 μM β-mercaptoethanol [gibco], 1000 units mL^-1^ mLIF). Every 3 days, the mESCs were passaged using Accutase.

mEpiSCs were cultured under 20% O_2_ and 5% CO_2_ at 37 °C condition. The plates should be coated by serum before use. mEpiSCs were maintained in FN medium (50% F12/Neurobasal [Gibco], N2 [Gibco, 200×] + B27 [Gibco, 200×], NEAA [Gibco, 100×], GlutaMAX [Gibco, 100×], BSA [1mg/mL], 100 μM β-mercaptoethanol [gibco], Activin A [20ng/mL], bFGF [20ng/mL]). Every 3 days, the mEpiSCs were passaged using Accutase.

### Generation of DsRed-labeled hPSCs and GFP-labeled mESCs

To construct UH10 hiPSCs with constitutive expression of DsRed in AAVS1 habor locus, guide RNA (gRNA) for AAVS1 safe harbor locus was designed on the website (crispr.mit.edu) and cloned into pX330 (a vector that can express Cas9 protein and guide RNA). Donor plasmid (pUC57-Neomycin-AAVS1-CAG-DsRed) contained left and right homology arms of AAVS1 safe harbor locus. 1 × 106 hPSCs were electroporated with 4 μg of donor plasmid and 2 μg of pX330 plasmid and DsRed+ cells were primarily selected by G418 (100 μg per mL) and then sorted by FACS (fluorescence-activated cell sorting). The GFP were cloned into a lentiviral vector Psin, and lentivirus was produced in 293T cells by cotransfecting. Viral supernatants were collected at 48 h after transfection and passed through a 0.45 µm filter to remove cell debris, then subjected to ultracentrifugation (20,000 × g for 3 h at 4 °C). mESCs were transduced with lentivirus and the positive cells were selected by puromycin (1 μg per mL).

### Generation of MYCN and BCl2 forced-expression hPSCs

The human MYCN and BCl2-2A-MYCN gene were cloned into a lentiviral vector tetO-FUW for tet-inducible expression of MYCN or Co-expression MYCN and BCl2. Lentivirus was produced in 293T cells by cotransfecting the tetO-FUW-MYCN or tetO-FUW-BCl2-2A-MYCN with three helper plasmids (pRSV-REV, pMDLg/pRRE, and vesicular stomatitis virus G protein expression vector), which provide the essential elements to package lentivirus. Viral supernatants were collected at 48 h after transfection and passed through a 0.45 µm filter to remove cell debris, then subjected to ultracentrifugation (20,000 × g for 3 h at 4 °C). DsRed+ hPSCs were transduced with lentivirus. The expression of MYCN and BCl2 are induced by exogenous addition of doxycycline (DOX) (2 μg per mL) and the positive cells were selected by puromycin (1 μg per mL).

### Co-differentiation of hPSCs and mESCs

hPSCs and mESCs were dissociated into single cells and counted. For mono-layer co-differentiation *in vitro*, 1×10^5^ hPSCs and 1×10^5^ mESCs were mixed and plated onto Matrigel-coated 6-well plate. The control group contained only 1×10^5^ hPSCs or mESCs. Cells are differentiated for 4 days in embryonic body (EB) differentiation medium (DMEM/F12 [Hyclone], 20% FBS, NEAA [1×, GIBCO], GlutaMAX [1×, GIBCO], 0.1% beta-Mercaptoethanol [GIBCO]). For mixed embryonic body (EB) co-differentiation *in vitro*, 1×10^5^ hPSCs and 1×10^5^ mESCs were mixed and cultured with EB differentiation medium in cell culture flask. For mixed teratoma formation *in vivo*, 1×10^6^ hPSCs and 1×10^6^ mESCs were mixed and resuspended in 30% matrigel (Corning) in DMEM/F12 (Hyclone), and then injected subcutaneously into NOD/SCID immunodeficient mice, obtained from Beijing Vital River Laboratory Animal Technology Co., Ltd. Teratomas were detected after 4 weeks. Mice were fed with the dox-containing water (2 mg/mL) for inducible MYCN, BCl2 or MYCN-BCL2 expression.

### Animal experiments

ICR mice were purchased from Beijing Vital River Laboratory Animal Technology Co., Ltd. *Flk-1*^*+/EGFP*^ mice were purchased from Jackson Lab (Bar Harbor, ME, USA). Female mouse at 4–6 weeks of age were selected as donors or surrogate mother. Two days before mating the donors were injected with 7.5 U pregnant mare’s serum gonadotropin (PMSG), and 48 h later the donors were injected with 7.5 U hCG and mated with the male mouse. The embryos at 8 cell stage or later morula to early blastocyst stage were obtained, respectively. Ten cells were injected into the embryos for following experiments. For in vitro chimerism assay, the embryo culture medium—KSOM (Millipore) was added with dox (2 μg per mL) for inducible MYCN or MYCN-BCl2 expression. The injected embryos were cultured in medium until blastocyst stage for further analysis. For in vivo chimera assay, the 10–20 embryos were transplanted into the uterus of per pseudopregnant mouse and the surrogate mice with embryos injected hPSCs were fed with the dox-containing water (2 mg per mL) until E10.5. The mice were euthanized, and embryos, vitelline membrane and placenta were obtained for subsequent analysis.

New Zealand rabbits were obtained from Huadong Xinhua experimental animal farms, District of Guangzhou. Female rabbits were injected with 100 IU PMSG, and 72-120 hours later further injected with 100 IU hCG and mated with the male rabbits. Embryos at 8 cell stage were harvested and injected with 7 indicated cells for in vitro assay, and cultured in EBSS (Thermo) Medium until blastocyst stage.

Pigs were purchased from a local slaughterhouse and cumulus oocyte complexes (COCs) were aspirated from antral follicles. After 44 hours in vitro maturation, the oocytes were activated with 2 successive DC pulses of 120 V per mm for 30 μseconds using an electrofusion instrument (CF-150B, BLS, Hungary). The activated oocytes were cultured in PZM-3 medium for partheno-development. After 48 hours or 4 days of culture, embryos at 8 cell stage were selected respectively for microinjection. 3-5 cells were injected to 8 cell stage embryos. The injected embryos were cultured in PZM3 medium (ENZO) until blastocyst stage for further analysis.

The interspecies chimerism experiments using hPSCs have been approved by the Ethnical Committee on Animal Experiments at Guangzhou Institutes of Biomedicine and Health, Chinese Academy of Sciences. Also, all the experiments were performed according to ISSCR guidelines.

### Chimerism analysis

hPSCs were completely dissociated using Accutase and centrifuged at 300 × g at room temperature for 3 min. After removal of the supernatant, cells were resuspended in the culture medium at a proper density (2–6 × 10^5^ cells per mL) and placed on ice for 20–30 min before injection. For in vitro extra-embryonic chimerism analysis, cells were microinjected into the eight-cell stage embryos. 10 indicated DsRed+ cells were injected in mouse embryos; 3-5 indicated DsRed+ cells were injected in pig embryos; 7 indicated DsRed+ cells were injected in rabbit embryos. While for in vivo chimerism analysis, 10 indicated DsRed+ cells were microinjected into the mouse later morulas or early blastocysts. For in vitro chimerism assay, the injected embryos at blastocyst stage were fixed by 4% PFA for 30 min and immunostained (CDX2, OCT4, and DAPI). For chimerism analysis in vivo, embryos were harvested until E10.5 for human mitochondria DNA assay, FACS or immunostaining.

### Single-cell cloning efficiency

For single-cell cloning assay of primed hPSCs, cells were plated onto 12-well plate at a density of 2000 cells per well, after dissociating into single cells with Accutase and centrifuged at 300 × g at room temperature for 3 min. Seven days later, cells were fixed in 4% PFA for 2 min. After washing with PBS, cells were stained with alkaline phosphatase staining solution (Beyotime) for 10−15 min.

### Immunofluorescence

The injected blastocyst embryos and frozen sections of E10.5 mouse chimeric embryos and placenta were incubated with antibodies (Stem121, CDX2, and OCT4) overnight at 4 °C, washed with PBS and incubated with specific secondary antibody. Nuclei were stained with DAPI. Stained embryos and sections were observed using a LSM800 confocal microscope (Carl Zeiss).

### Flow cytometry and cell sorting

The embryonic tissue were digested with collagenase and washed with PBS, and directly detected by flow cytometry. We randomly selected and pooled 10 mouse embryos together to sort the cells. The embryonic tissues were digested with collagenase and washed with PBS, centrifuged to remove the supernatant, then washed with PBS and stained with human cells specific antibody TRA-1-85 and human CD34 antibody (direct-labeled antibody) at 4°C for 30 minutes, finally, the live TRA-1-85^+^DsRed^+^ cells were obtained by flow cytometry sorting.

### CFU assay and Cell morphology

The CFU assay was performed according to the manufacturer’s instruction of Methocult H4435 (Stem Cell Technologies). Firstly, cells were suspended into 120 μl IMDM medium supplemented with 2% FBS (Biological Industries), and then add the cell suspension to 1 ml Methocult H4435. Next, transferred the mixture to 35 mm ultra-low attachment plates (Stem Cell Technologies) and rotated gently to spread methylcellulose medium over the surface of the dish. Placed 3 dishes within a 100 mm petri dish containing with 3 mL sterile water and incubated the dishes in 37 °C, 20% O_2_ and 5% CO_2_ condition. The CFUs were classified and calculated according to the morphology after 2 weeks. Then, harvested cells from the CFU assay, rinsed twice with DPBS (gibco) and resuspended the cells in 200 μl DPBS. The morphology of cells was showed by microscopy after cytospin (500g, 3 min; cence) and stained with May–Grunwald–Giemsa.

### Western blot assays

Cells were lysed on ice with 200 µL of RIPA buffer (Beyotime) for 15 min and separated by 12% sodium dodecyl sulfate–polyacrylamide gel electrophoresis (SDS-PAGE) before being transferred onto polyvinylidene difluoride (PVDF) membranes (Millipore). The membranes were blocked in 5% nonfat milk for 1 h and incubated overnight at 4 °C with the appropriate diluted primary antibodies or anti-flag/GAPDH antibody. Subsequently, the membranes were incubated with HRP-conjugated secondary antibody for 2 h at room temperature and HRP was detected by ECL (Advanste) and visualized by SmatChemi Image Analysis System (SAGECREATION).

### Quantitative real-time PCR

Total RNA was extracted with the RaPure Total RNA Micro Kit (Magen). Two-microgram RNA was reversing transcribed into cDNA and amplified with SYBR Green PCR Master Mix (Bio-Rad).

### Teratoma formation

The teratoma formation experiments were approved by the Ethical Committee on Animal Experiments at Guangzhou Institutes of Biomedicine and Health, Chinese Academy of Sciences. 1×10^6^-6×10^6^ Cells were digested by Accuatse (Sigma) for 5 min at 37 °C and resuspended in 30% matrigel (Corning) in DMEM/F12 (Hyclone), and then injected subcutaneously into NOD/SCID immunodeficient mice, obtained from Beijing Vital River Laboratory Animal Technology Co., Ltd. Teratomas were detected after 4 weeks and fixed in 4% PFA. After paraffin embedding and sectioning, sections were stained with hematoxylin/eosin.

### Histopathological observation

After formalin fixation, embryo samples were embedded in paraffin, sectioned at 3-mm thickness, and stained with hematoxylin and eosin (H&E) for histopathological examination.

### Immunohistochemistry

Embryo tissue samples were fixed in 4% formalin, embedded in paraffin and sectioned at 3μm thickness. Embryo tissue samples were fixed in 4% formalin, embedded in paraffin and sectioned at 3μm thickness. Lung sections were treated with xylene to remove the paraffin and then were rehydrated. Prior to staining heat-induced antigen retrieval was performed by placing the slides into 0.01M citrate buffer solution (pH6.0), and subjected to microwave heating three times for 5min. Then the sections were incubated with 3% H2O2 for 10min at room temperature and washed three times with PBS, followed by incubation with serum for 30min. Samples were incubated with the antibodies (Stem121) overnight at 4°C. After washing with PBS, the slices were incubated with secondary antibodies for 0.5h at 37°C. After staining with 3, 3’-diaminobenzidine (DAB), the sections were observed under optical microscope.

### Statistics

In general, data were presented as mean + SEM, and statistics were determined by unpaired two-tailed Student’s test (t test). P value < 0.05 was considered statistically significant. *p < 0.05; **p < 0.01; ***p < 0.001. No statistical method was used to predetermine the sample size. No samples were excluded for any analysis. No randomization was used for allocating animal group. No blinding done in animal experiments.

## Data availability

The datasets generated and/or analysed during the current study are available from the corresponding author on reasonable request.

## Figure legends

**Supplemental figure 1**

**(A)** The morphology of MYCN or MYCN/BCL2 expressed hPSCs. Scale bars, 100 µm. **(B)** RT-qPCR analysis of pluripotency markers and other lineage genes in MYCN or MYCN/BCL2 expressed hPSCs. **(C)** Flow cytometry analysis of the pluripotency markers in MYCN-hPSCs (M-hPSCs) and MYCN/BCL2-hPSCs (M/B-hPSCs). **(D)** The cell growth rate of hPSCs, M-hPSCs and M/B-hPSCs under normal maintenance culture.**(E)** Karyotype analysis of M-hPSCs and M/B-hPSCs. **(F)** Hematoxylin and eosin (H&E) staining of the teratoma formed by M-hPSCs in the presence or absence of Dox (2mg/ml). Scale bars, 100 µm.

**Supplemental figure 2**

**(A)** Schematic overview of co-differentiation of DsRed labeled hPSCs with mESCs *in vitro*. hPSCs-DsRed and mESCs-GFP are initially mixed in 1:1 ratio at single cell for further differentiation. **(B-C)** Analysis of survival of hPSCs-DsRed or mESCs-GFP in co-differentiation in vitro. Dissociated hPSCs-DsRed or mESCs-GFP cells are cultured in differentiation condition either alone or mixed with each other in 1:1 ratio for 4 days. Scale bars, 50 µm. **(D)** Flow cytometric analysis of Annexin V^+^ human cells from hPSC+mESC co-differentiation condition for 4 days. Error bars represent mean+SEM of three independent replicates. ***p < 0.001.

**Supplemental figure 3**

**(A)** Schematic overview of co-differentiation of DsRed labeled hPSCs or MYCN-hPSCs with mESCs *in vitro*. hPSCs-DsRed and mESCs-GFP are initially mixed in 1:1 ratio at single cell for further differentiation. **(B-D)** Analysis of human cell proliferation rate and Annexin V^+^ cells on the mixture of hPSCs-DsRed or MYCN-hPSCs-DsRed and mESCs cultured in differentiation condition for 4 days. Error bars represent mean+SEM of three independent replicates. ***p < 0.001. **(E)** Cloning efficiency is analyzed by alkaline phosphatase staining on colonies formed by the individualized cell of the indicated cell lines. 2000 individualized were plated in each well of 12-well-plate and cultured for 7 days. **(F)** Analysis the survival of Dsred labeled hPSCs in pre-implantation pig embryos. DsRed^+^ hPSCs (3-5 cells for pig) were injected into 8-cell (8C) stage embryos and cultured for 3 days. Scale bars, 20 µm.

**Supplemental figure 4**

**(A)** Analysis the cell growth rate and apoptosis rate of hPSCs or M/B-hPSCs during co-differented with primed mPSCs (mEpiSCs) in 1:1 ratio for 4 days. **(D)** Analysis the cell counts and apoptosis rate of hPSCs or M/B-hPSCs during co-differented with mESCs in different ratio. **(C)** Teratomas formed by M/B-hPSCs mixed with mESCs in different ratio. **(D)** Analyses of human cells at different embryonic regions in E10.5 mouse chimera by HE staining and Immunohistochemical staining using human specific marker Stem121. Scale bars, 500 µm (D-left), 20 µm (D-middle), 5 µm (D-right). **(E)** Representative placenta confocal images showing DsRed^+^ human cells can contribute to trophoblastic lineages in chimeric E10.5 placentas. The placentas were stained with anti-CK7 (trophoblastic lineages marker). Scale bars, 20 µm. **(F)** Co-differentaiton assay mixed another M/B-hPSCs cell line (HN10+N+B) with mESCs in 1:1 ratio for 4 days.

