## Supplementary figures and images for "Generating Functional Cells Through Enhanced Interspecies Chimerism with Human Pluripotent Stem Cells"

### Supplemental figure 1

##### Supplemental figure 1

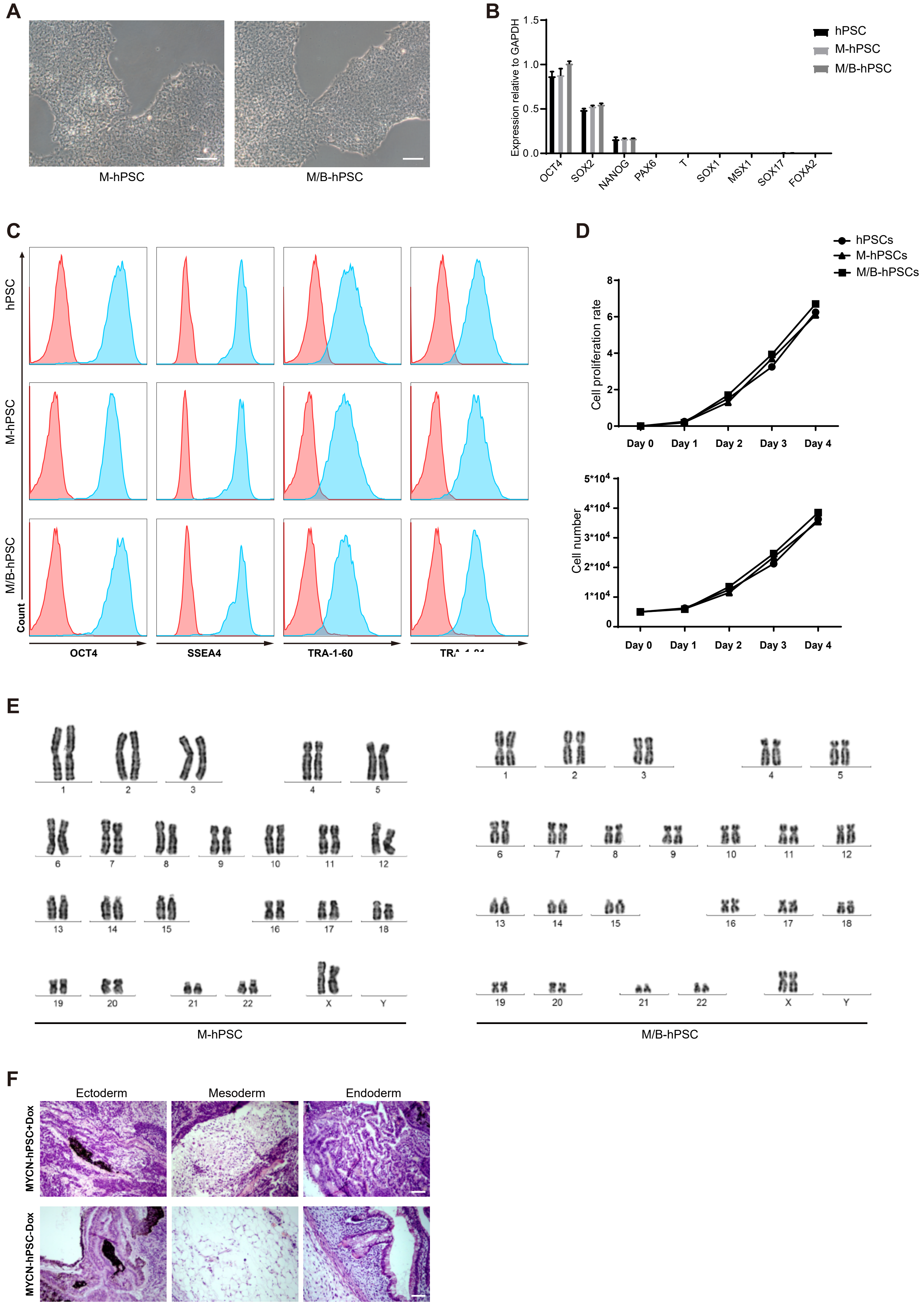

### Supplemental figure 2

##### Supplemental figure 2

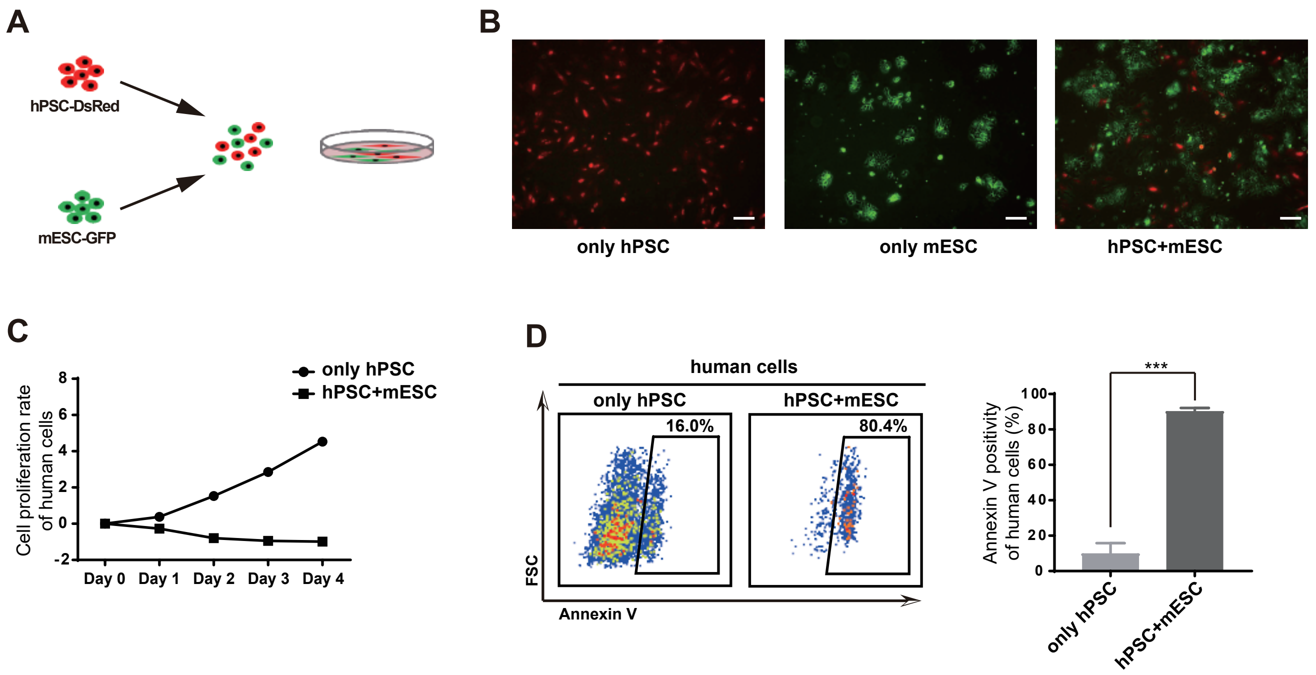

### Supplemental figure 3

#### Supplemental figure 3

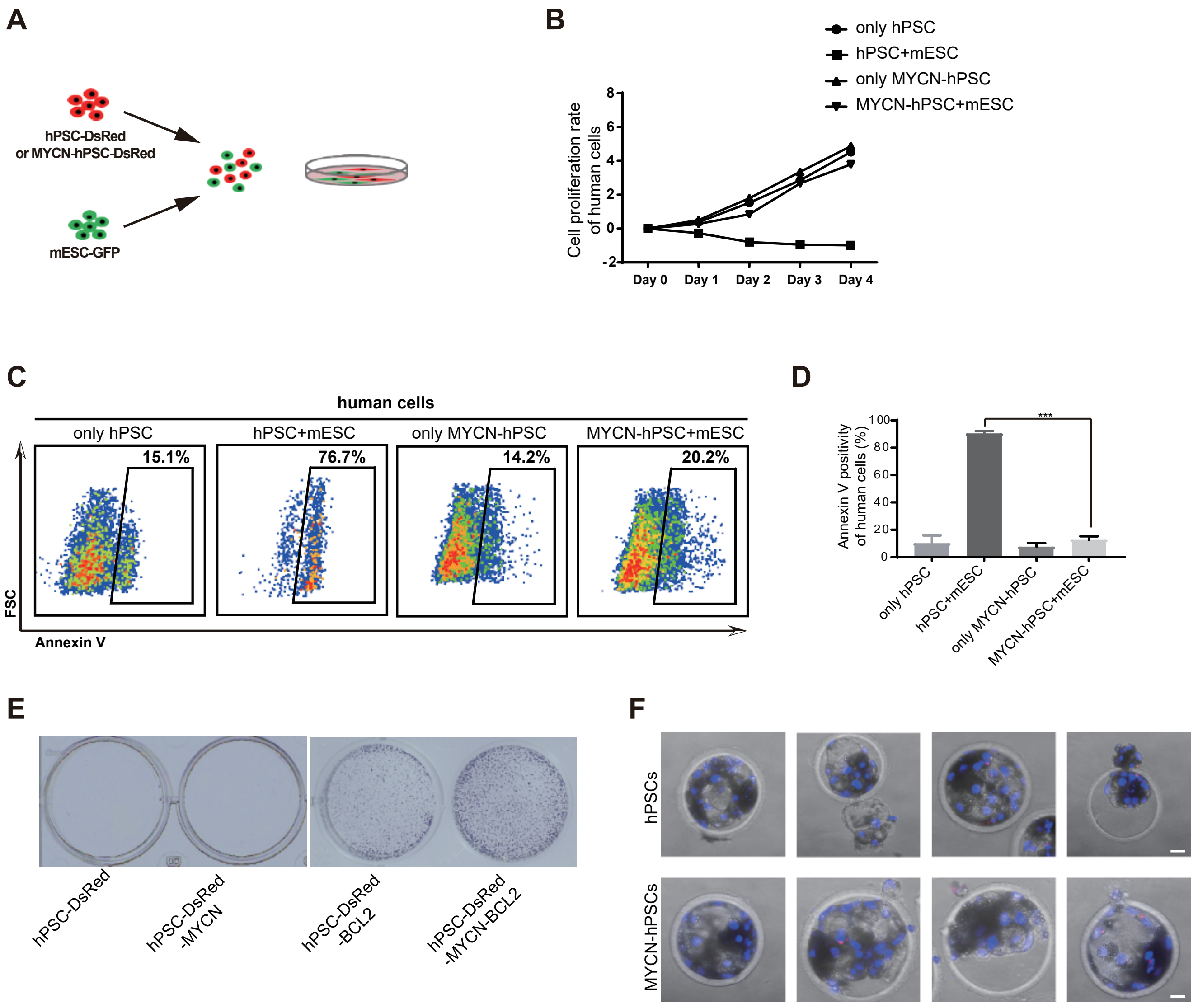

### Supplemental figure 4

#### Supplemental figure 4

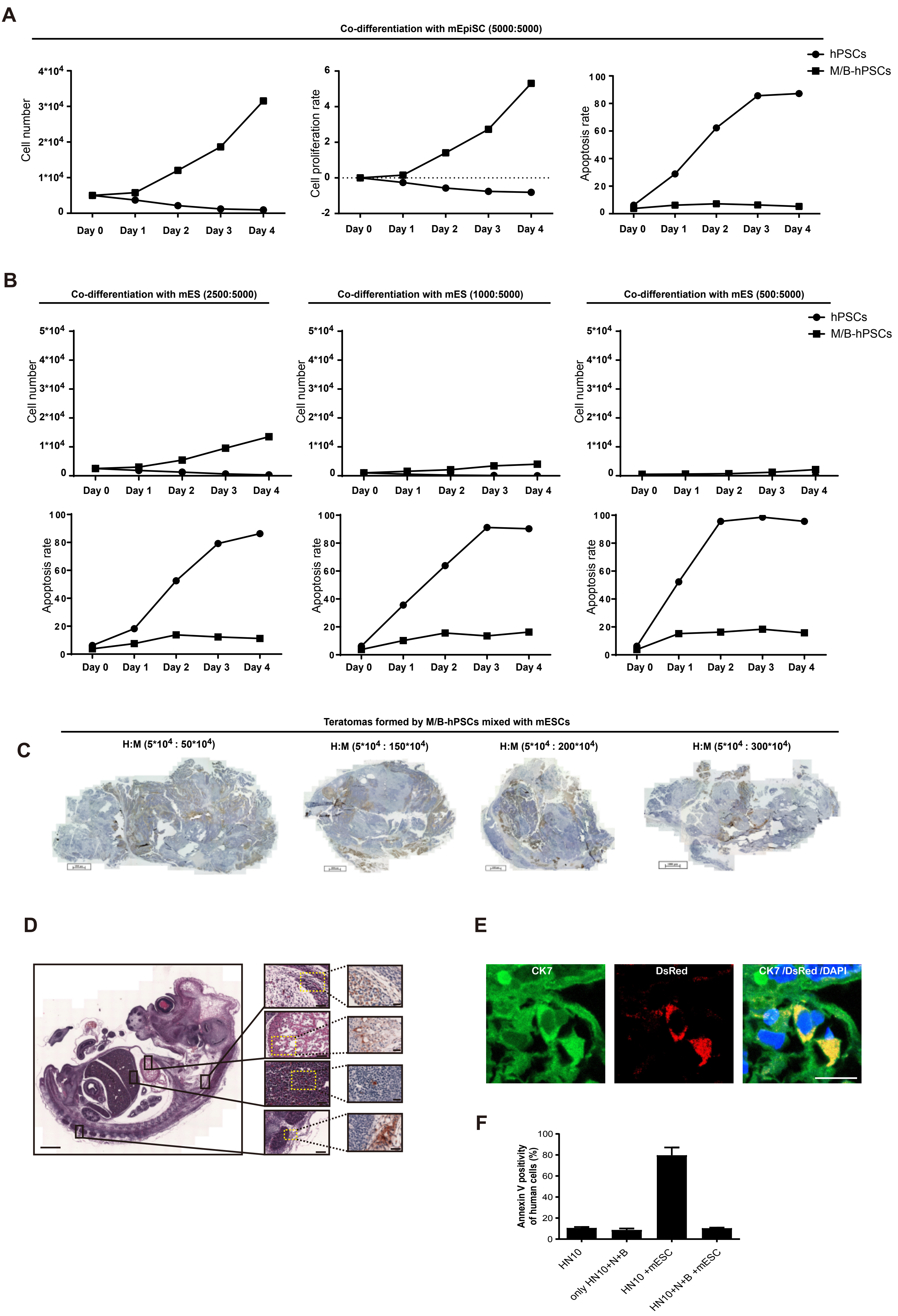
