## Supplemental Table 1 for "Generating Functional Cells Through Enhanced Interspecies Chimerism with Human Pluripotent Stem Cells"

**Supplemental Table 1 primers used in qRT-PCR**

| | |
|---|---|
| Q-H-ß- actin-F | CT CCAT CCT GGCCT CGCT GT |
| Q-H-ß- actin -R | GCT GT CACCT T CACCGT T CC |
| Q-M-ß- actin -F | CGT T GACAT CCGT AAAGACC |
| Q-M-ß- actin -R | AACAGT CCGCCT AGAAGCAC |
| Q-H-MYCN-F | ACCCGGACGAAGAT GACT T CT |
| Q-H-MYCN-R | CAGCT CGT T CT CAAGCAGCAT |
| Q-M-MYCN-F | CCT CACT CCT AAT CCGGT CAT |
| Q-M-MYCN-R | GTGCTGT AGT T T T TCGT TCACTG |
| Q-H-OCT4-F | CCT CACT T CACT GCACT GT A |
| Q-H-OCT4-R | CAGGT T T T CT T T CCCT AGCT |
| Q-H-SOX2-F | CCCAGCAGACT T CACAT GT |
| Q-H-SOX2-R | CCT CCCAT T T CCCT CGT T T T |
| Q-H-NANOG-F | T GAACCT CAGCT ACAAACAG |
| Q-H-NANOG-R | T GGT GGT AGGAAGAGT AAAG |
| Q-H-PAX6-F | AT GT GT GAGT AAAAT T CT GGGCA |
| Q-H-PAX6-R | GCT T ACAACT T CT GGAGT CGCT A |
| Q-H-T-F | T AT GAGCCT CGAAT CCACAT AGT |
| Q-H-T-R | CCT CGT T CT GAT AAGCAGT CAC |
| Q-H-MSX1-F | T CCGCAAACACAAGACGA |
| Q-H-MSX1-R | ACT GCT T CT GGCGGAACT T |
| Q-H-SOX17-F | CGCACGGAAT T T GAACAGT A |
| Q-H-SOX17-R | GGAT CAGGGACCT GT CACAC |
| Q-H-FOXA2-F | ACT ACCCCGGCT ACGGT T C |
| Q-H-FOXA2-R | AGGCCCGT T T T GT T CGT GA |
| Q-H-hmtDNA-F | AAT AT T AAACACAAACT ACCACCT ACCT |
| Q-H-hmtDNA-R | T GGT T CT CAGGGT T T GT T AT AA |
| Q-H-UCNE-F | AACAAT GGGT T CAGCT GCT T |
| Q-H-UCNE-R | CCCAGGCGT AT T T T T GT T CT |
| Q-H-MYC-F | GGCTCCTGGCAAAAGGTCA |
| Q-H-MYC-R | CTGCGTAGTTGTGCTGATGT |
