## Supplemental Table 2 for "Generating Functional Cells Through Enhanced Interspecies Chimerism with Human Pluripotent Stem Cells"

**Supplemental Table 2 Antibodies**

| Antibodies | Vendor | Cat# | Dilution |
|---|---|---|---|
| Anti-human CD34- PerCP-Cy5.5 | BD Biosciences | 347203 | 1:100 |
| Anti-human OCT4 | BD biosciences | 560307 | 1:100 |
| Anti-human SSEA4 | Invitrogen | 414000 | 1:100 |
| Anti-human TRA-1-60 | Santa Cruz Biotechnology | sc-21,705 | 1:100 |
| Anti-human TRA-1-85-APC | RD | FAB3195A | 1:100 |
| Anti-human TRA-1-81 | Santa Cruz Biotechnology | sc-21,706 | 1:100 |
| Anti-human CDX2 | Thermo | MA5-14494 | 1:100 |
| Anti-human stem121 | TAKARA | Y40410 | 1:100 |
| Anti-human CD45 | BD Biosciences | 560973 | 1:100 |
| DAPI | Thermo | 62248 | 1:5000 |
